# Digital imaging outperforms traditional scoring methods of spittlebug tolerance in *Urochloa humidicola* hybrids

**DOI:** 10.1101/2020.10.13.338186

**Authors:** Luis M. Hernandez, Paula Espitia, Valheria Castiblanco, Juan A Cardoso

**Author notes:** Correspondence to; Tropical Forages Program. Alliance for Bioversity-CIAT. Regional Hub for the Americas. A.A, 6713, Cali, Colombia.

## Abstract

American spittlebug complex (Hemiptera: Cercopidae) is a critical pest for existing *Urochloa humidicola* cultivars in the neotropical savannas. The *U. humidicola* breeding program of the International Center for Tropical Agriculture aims to increase tolerance to spittlebugs. To develop *U. humidicola* genotypes with superior tolerance to spittlebugs than existing cultivars, adequate screening methods ought to be deployed. Currently, visual scores of plant damage by spittlebugs is the standard method to screen for variation in plant tolerance. However, visual scoring is prone to human bias, is of medium throughput and relies of the expertise of well-trained personnel. In this study, we compared estimations of plant damage from two alternative methods (SPAD measurements and digital images) and visual scoring from an inexpert evaluator with the plant damage estimated from an expert. This information should instruct if different methods could be implemented in the *U. humidicola* breeding program. Time needed to evaluate damage was recorded for each method. Lin’s correlation coefficient, Pearson’s correlation coefficient and broad sense heritability values were also calculated. Overall, damage estimated from digital images showed the highest throughput (twice as fast as visual scoring from an expert); high correlations with visual scoring (*r* > 0.80, p < 0.0001); and heritability values for plant damage as good or better (> 0.7) than those obtained by visual scoring from an expert. Our results indicate that digital imaging is a phenotyping method that might improve the efficiency of breeding for increased tolerance to spittlebugs in *U. humidicola*.

**Highlight:** Digital imaging outperformed standard scoring method of spittlebug tolerance in *Urochloa humidicola,* suggesting that this method might improve the efficiency of breeding for such stress.

## Introduction

*Urochloa humidicola* is an important forage grass in the tropical savannas of America (Vasques Berchembrock et al. 2020). The productivity of current cultivars of *U. humidicola* is challenged by the American spittlebugs complex (Valério et al. 2001). Increasing tolerance to spittlebugs complex in *U. humidicola* is a major target for the *Urochloa* breeding program of the International Center for Tropical Agriculture (CIAT, Colombia). To develop *U. humidicola* genotypes with superior tolerance to spittlebugs than current available cultivars, adequate screening methods ought to be deployed. Currently, visual scoring of plant damage is the standard phenotyping method to evaluate plant tolerance to spittlebug complex in *Urochloa* grasses. Visual scores rely on estimates of percentages of dead leaf tissue (Parsa et al. 2011). Overall, visual scoring is a low cost and medium throughput phenotyping method that has proven successful in the *Urochloa* breeding program of CIAT (Cardona et al. 1999; Miles et al. 2006).

Visual scoring is nonetheless prone to the subjectivity of any given evaluator and may not be accurate enough (Walter et al. 2012). Among factors that can affect scoring of plants are the expertise of the evaluator (i.e., different scores from different evaluators) and fatigue over working hours. To overcome this, sensor based measurements are gaining momentum in the *Urochloa* breeding program (Cardoso & Rao 2019). Hand-held devices such as the SPAD series meters are used to non-destructively record greenness of leaves. Measurements of SPAD have been shown to be positively and linearly correlated with percentages of dead tissue in *Urochloa* grasses (Cardoso et al. 2013). Another method used to record percentages of dead leaf tissue in *Urochloa* grasses is digital imaging (Jiménez et al. 2017).

Albeit sensor based measurements are currently used in the *Urochloa* breeding program, their use have been limited to evaluations different to those of tolerance to spittlebugs (e.g., Jiménez et al. 2017; Cardoso & Rao 2019; Mazabel et al. 2020; Jiménez et al. 2020). Therefore, the main objective of the preset work was to compare how well alternative phenotyping methods (SPAD measurements and digital images) or a visual scoring from an inexperienced evaluator related to the traditional evaluation based on visual scoring of damage from an expert. For that purpose, a set of 24 *U. humidicola* hybrids (plus seven *U. humidicola* genotypes with known tolerance to spittlebugs) were used and evaluated under greenhouse conditions. Estimations of plant damage using different methods were carried out and calculations of agreement (Lin’s concordance index) and Pearson correlation coefficient were performed. Broad sense heritability value was calculated to provide an indication of the efficiency of the selection process from the different phenotyping methods. This information should instruct which screening methodology is the most appropriate (in terms of ease, accuracy and throughput), but also guide further refinements needed in any screening method used. Improved screening methods should allow more accurate and intense selection, and hence, greater genetic gain for tolerance to spittlebugs in *U. humidicola* hybrids.

## Materials and Methods

Thirty-one *U. humidicola* genotypes were used in the present study, which was conducted at CIAT (Palmira, Colombia, latitude 3°31’N; longitude 76°19’W; altitude 965 m). Genotypes included 24 hybrids originated from the *U. humidicola* breeding Program of CIAT, and seven plant checks with known tolerance to spittlebugs. Plant checks consisted of three tolerant genotypes (cvv. Llanero and Tully and one germplasm accession, CIAT/16888) and three sensitive ones (two germplasm accessions, CIAT/26146, CIAT/26375 and a hybrid, Bh13/2768). The germplasm accessions CIAT/16888 and CIAT/26146 are the foundation parents of the *U. humidicola* breeding program. All genotypes were obtained from vegetative material (propagation plants) that were maintained under greenhouse conditions [28°C; 80% RH]. For each genotype, ten vegetative propagules (plant units) of one single tiller were harvested from propagation plants and then immersed for five minutes in a 1% sodium hypochlorite solution. Plants were rinsed from sodium hypochlorite prior to planting. Each plant unit was planted in a cylindrical polyvinyl chloride (PVC) unit (5.3 cm wide X 6.5 cm deep) that contained 40 g of sterilized soil (3:1 weight soil: weight sand). Plants were watered daily and fertilized with 30 mL of nutrient solution prepared with a 15% N-15% P-15% K soluble fertilizer at 3 g L^-1^ two weeks after planting. One month after planting, when sufficient superficial roots were available to serve as feeding sites for the nymphs, five plants/genotype were infested with six mature eggs of *Aeneolamia varia* as previously described by (Cardona et al. 1999). The other five plants/genotype were not infested and used as controls. The eggs were previously obtained from the CIAT spittlebug mass rearing colony, selected for viability by visual inspection and incubated under controlled conditions (28C, 85% RH) (Parsa et al. 2011). Plants were organized in a randomized complete block with two treatments (infested with *A. varia* and uninfested) and five replicates.

### Plant Damage evaluation

Three phenotyping methods for plant damage were carried out at weekly intervals for five weeks: 1) visual scoring from an expert and an inexpert evaluator; 2) SPAD measurements and, 3) digital images. Plant damage was estimated and expressed in percentage as described below. Also, time spent during plant damage evaluation under the different methods was recorded.

### Visual scoring

Visual scoring for plant damage consisted on the assessment of the proportion of green to senescing leaf tissue (yellow to brown) of the whole plant. Visual scoring used a 11-point scale as follows: 0 = all leaves are green; 1= 10% of senescent leaves; 2 = 20% of senescent leaves; …..; 10 = 100% of senescent leaves. To test whether the visual scoring was affected by a given person during an evaluation, an expert and an inexpert evaluator carried out visual scorings independently.

### SPAD measurements

SPAD meters (SPAD-502, Konica Minolta, Japan) were used to estimate greenness of different leaves. SPAD units were recorded on three fully expanded leaves for each plant and their values averaged. Plant damage was estimated from the difference in SPAD measurements between consecutive weeks as follows; [(SPAD_n_-SPAD_n+1_)/SPAD_n_]*100. were SPAD_n_ is a SPAD recording at any given week, and SPAD_n+1_, the recording of SPAD the week after.

### Digital imaging

For image acquisition, individual plants were placed within a closed chamber (dimensions: 2×1.5×1 m) and illuminated from above with a T8 led tube 32w 120 cm. Images were then acquired with a digital color camera (Nikon Coolpix P6000, Nikon, Japan) with the following set up: F-stop: f/2.7, Exposure time: 1/60, and ISO speed ISO-89 and from a Nadir view of the plant. Images were saved in a 4224 x 3168 pixel JPEG format. To account for difference in illumination throughout acquisitions and thereby influencing color tones in images and greenness estimates, images were pre-processed with GIMP software (GIMP 2.10). GIMP software was used to apply a pre-saved color tone matching curve to all JPEG files. After that, images were processed and analyzed using ImageJ (ImageJ 1.51). Image processing consisted on splitting the images onto their color channels (Red, Green and Blue), and then normalizing the blue channel (Blue channel / Red channel + Green channel + Blue channel). The normalized blue channel was used for image segmentation using the default threshold method of ImageJ. Image segmentation consisted on the separation of shoot (white pixels) from background (black pixels). Once the image was segmented, a mask was laid onto the original unsegmented image using the AND logic operation. The masked image was then used to calculate the difference between green and red channels, which enhances contrast between green tissue and senescing (yellow to brown) tissues. Once the normalized green red difference index was calculated, K-means clustering was used to create three clusters of colors in the image: background, green tissue and senescing (yellow to brown) tissue. The number of pixels for each cluster was then quantified and plant damage was calculated as ([SP/(SP+GP)]*100; where SP = number of pixels clustered as senescing tissue and GP = number of pixels clustered as green tissue. Figure 1. summarizes the image processing pipeline.

**Figure 1.**
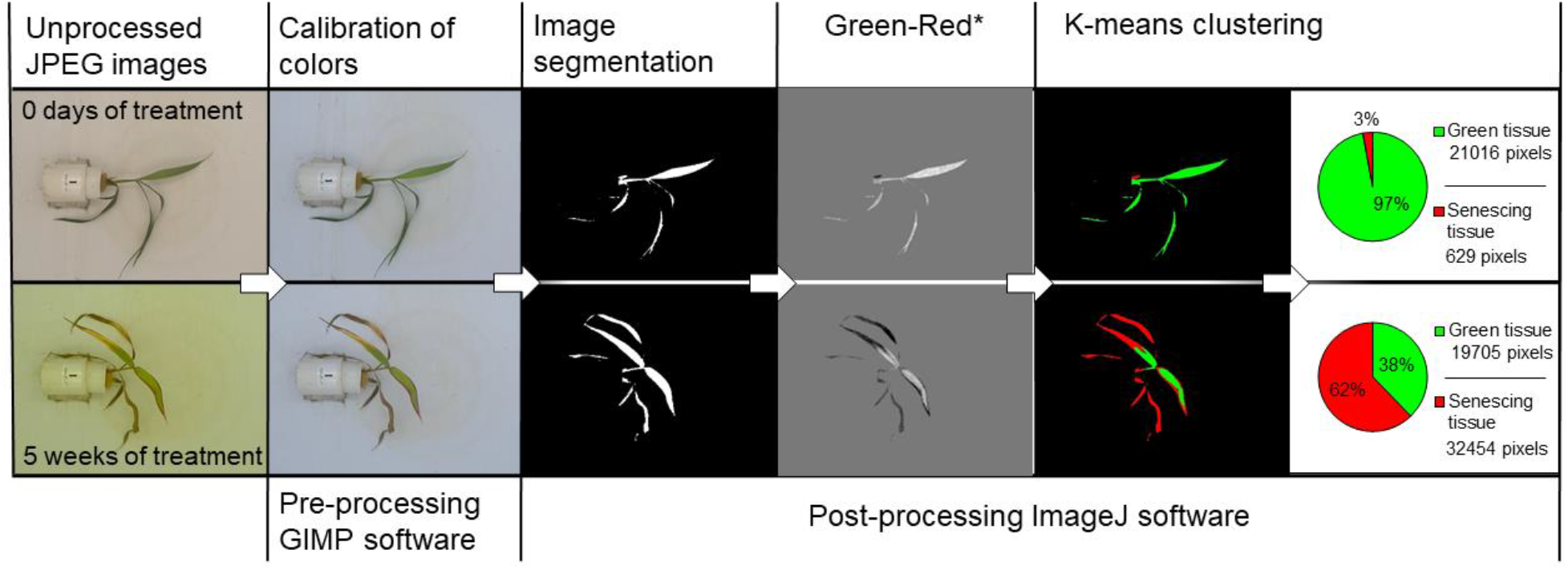
Summary of the image processing pipeline. Green-Red* is the product of the green minus red channel.

### Statistical analysis

Mean values and standard deviations were calculated for estimations of plant damage for different dates and evaluation methods. Two-way analyses of variance were calculated. Analyses were performed only in infested plants and conducted in R (R Development CoreTeam 2015). Calculations of agreement (Lin’s concordance index; Lin, 1989) and Pearson correlation coefficient were performed between estimates of plant damage from alternative methods (SPAD meter and digital imaging) and visual scores from an inexpert evaluator with the standard evaluation of visual scoring of plant damage from an expert. Broad sense heritability was calculated for each of the different evaluation methods (Piepho & Möhring 2007).

## Results and discussion

### Comparison of throughput and estimated damage from phenotyping methods

The present study aimed to compare how well alternative phenotyping methods (SPAD meter and digital images) or a visual scoring from an inexperienced evaluator related to the standard evaluation of visual scoring of plant damage from an expert. Among them, the throughput of each method (in terms of time consumed by one person to evaluate plants) was compared (Table 1). Overall, capture of digital images was the fastest method to record plant damage (twice as fast as second fastest, i.e., visual scoring from expert evaluator). Similar results were previously shown by several authors (Büchi et al. 2018; Jiménez et al. 2020). Reduction of time is among the improvements sought by most phenotyping methods (c.f., Shakoor et al. 2017; Araus et al. 2018). Faster phenotyping could allow the increment of number of plants to be evaluated for plant damage and/or to reduce the time dedicated to such activity; or allow more intensive phenotyping (recording of additional traits that might be of interest).

**Table 1.** Average values of five evaluations showing the time required to perform evaluations. Values denote means ± standard deviations. Different letters next to standard deviation values denote significant differences at α = 0.05.

|  | Number of plants/hour |
| --- | --- |
| Visual scoring (expert) | 58 ± 13b |
| Visual scoring (inexpert) | 71 ± 25c |
| SPAD measurements | 80 ± 15bc |
| Digital images | 113 ± 15a |

Estimations of plant damage using different phenotyping methods are shown in Figure 2. Our results showed that there were not significant differences between estimates of damage from visual scoring from an expert and an inexpert evaluator. This suggests that the inexpert evaluator followed well instructions given by the expert evaluator. However, this might not always be the case for every training of new evaluators. The success of training of a new evaluator is dependent to inherent characteristics of such individual (e.g., previous knowledge of the plants; this case), which likely affects the accuracy of any evaluation. Bock et al. (2020) recently reviewed inter-rater variability and success of training among drawbacks of visual estimates of plant damage. Albeit estimates of damage from the expert and inexpert evaluators were similar, measures of data variability (i.e., standard deviation) from the inexpert evaluator were greater than those from the expert evaluator. Similar results were found by (El Jarroudi et al. 2015) when comparing estimates of septoria leaf blotch severity (and measures of data variability) in winter wheat from different evaluators.

**Figure 2.**
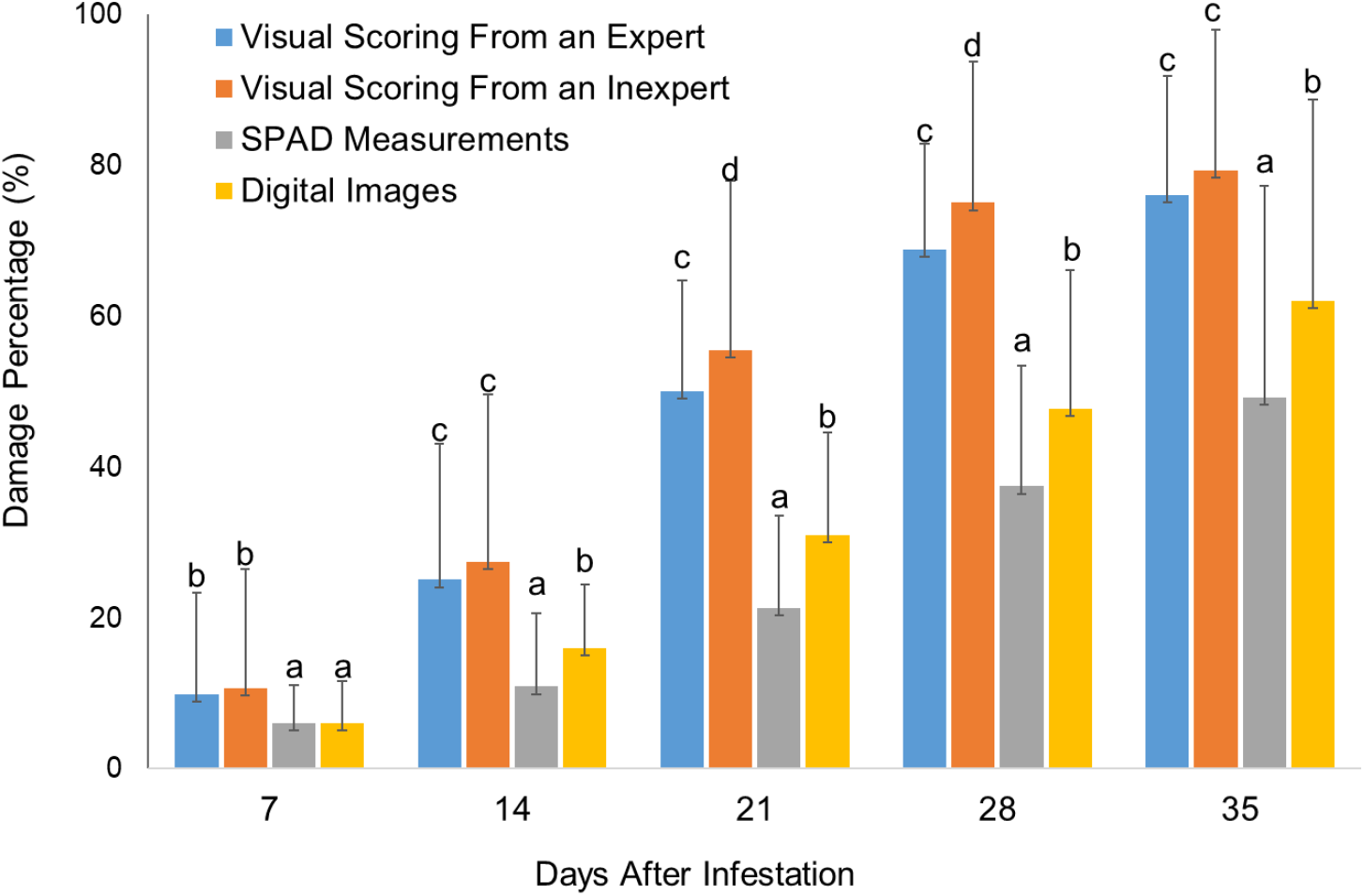
Comparison of damage percentage of three plant damage evaluation methods assessed over Bh16 genotypes at 7, 14, 21, 28, and 35 days after infestation with spittlebug nymphs *Aeneolamia varia.* Letter over bars indicated the differences by evaluation method at 7, 14, 21, 28 or 35 days after infestation. Column bars represent means and error bars indicate the standard deviation. Columns with different letter are significantly different (α < 0.05).

Differences in estimates of damage between visual scores (from expert and inexpert evaluators) and the other two methods (SPAD measurements and digital images) were found from the first week of evaluation (Table 2). Throughout the experiment, it was notable that estimates of damage were greater in visual scores compared to those obtained from SPAD meters (~ 1.5-fold greater) and digital images (~1.3-fold greater). Furthermore, development of damage appeared faster under the visual scoring method. Since the magnitude and speed of damage were greater under the visual scoring method (for both experienced and inexperienced evaluators), it is likely that visual scores over-estimated damage. Over-estimation of damage by visual scoring compared to other methods, including digital imaging, was previously identified by (Bock et al. 2010).

**Table 2.** Results of analysis of variance (two-way ANOVA) for 27 genotypes from *Urochloa humidicola* Bh16 population breeding program evaluated against spittlebug nymph *Aeneolamia varia*.

| DAI<sup>†</sup> | Source | Df | Sum of Square | Mean of Square | F value | P value |  |
| --- | --- | --- | --- | --- | --- | --- | --- |
| 7 | Genotype | 29 | 13869 | 478.2 | 4.849 | 7.06E-14 | *** |
|  | Method | 3 | 2471 | 823.7 | 8.352 | 0.000021 | *** |
|  | Genotype:Method | 87 | 9659 | 111 | 1.126 | 0.225 |  |
|  | Residuals | 416 | 41026 | 98.6 |  |  |  |
| 14 | Genotype | 29 | 33021 | 1139 | 5.712 | <2e-16 | *** |
|  | Method | 3 | 24089 | 8030 | 40.276 | <2e-16 | *** |
|  | Genotype:Method | 87 | 15286 | 176 | 0.881 | 0.761 |  |
|  | Residuals | 416 | 82935 | 199 |  |  |  |
| 21 | Genotype | 29 | 48623 | 1677 | 8.695 | <2e-16 | *** |
|  | Method | 3 | 103661 | 34554 | 179.185 | <2e-16 | *** |
|  | Genotype:Method | 87 | 12537 | 144 | 0.747 | 0.951 |  |
|  | Residuals | 416 | 80220 | 193 |  |  |  |
| 28 | Genotype | 29 | 64296 | 2217 | 11.771 | <2e-16 | *** |
|  | Method | 3 | 124900 | 41633 | 221.037 | <2e-16 | *** |
|  | Genotype:Method | 87 | 9217 | 106 | 0.562 | 0.999 |  |
|  | Residuals | 416 | 78355 | 188 |  |  |  |
| 35 | Genotype | 29 | 121290 | 4182 | 12.627 | <2e-16 | *** |
|  | Method | 3 | 76836 | 25612 | 77.323 | <2e-16 | *** |
|  | Genotype:Method | 87 | 20665 | 238 | 0.717 | 0.97 |  |
|  | Residuals | 416 | 137794 | 331 |  |  |  |

\*\*\*Significant at the 0.001 probability level.
<sup>†</sup>DAI, days after infestation.

### Concordances, correlations, and heritability

Highest concordances (Lin’s concordance coefficient, CCC) and correlations (*r*) were observed between visual scoring from the expert and inexpert evaluators (Table 2). Our results showed increasing values of CCC and *r* between visual scores from an expert and inexpert evaluator with each passing week until the end of the evaluation period (data not shown). This suggests that the inexpert evaluator got better with time in the visual scoring of plant damage, as shown elsewhere (Bock et al. 2016; Bock et al. 2020). Albeit the improvement gained by the inexpert evaluator, this one was not able to distinguish percentages of damage below 20% intervals, whereas the expert evaluator could do it at 10% intervals (data not shown). Similar results were found when experienced and inexperienced evaluators assessed severity of Phomopsis leaf blight of strawberry (Nita et al. 2003). Second best values of CCC (0.82) and *r* (0.87) were between visual scoring from expert evaluator and digital images (Table 3). Such level of agreement between estimates of damage from visual scoring and digital images is considered low (c.f., McBride, 1985). This is not surprising as 1) estimates of damage from visual scoring were discrete values vs. continuous values of plant damage estimated from digital images (c.f., McBride, 1985); and 2) a likely overestimation of damage from visual scoring (Figure 2) as mentioned before.

**Table 3.** Lin’s concordance coefficient and Pearson correlation analysis between damage percentages obtained from the evaluation methods. CCC: Lin’s concordance correlation coefficient, CI: confidence interval (95%), *r* = Pearson correlation coefficient, *** correlation significance (*p* < 0.001).

| DAI* | Index | Visual Scoring<br>From an Expert | Visual Scoring<br>From an Expert | Visual Scoring<br>From an Expert |
| --- | --- | --- | --- | --- |
|  |  | vs.<br>Visual Scoring<br>From an Inexpert | vs.<br>SPAD<br>Measurements | vs.<br>Digital Images |
| 7 | CCC | 0.89 | 0.16 | 0.29 |
|  | CI | 0.86 - 0.91 | 0.09 - 0.24 | 0.22 - 0.36 |
| | $r$ | 0.9*** | 0.25*** | 0.44*** |
| 14 | CCC | 0.89 | 0.24 | 0.42 |
|  | CI | 0.86 - 0.91 | 0.17 - 0.32 | 0.36 - 0.49 |
| | $r$ | 0.91*** | 0.36*** | 0.59*** |
| 21 | CCC | 0.89 | 0.34 | 0.58 |
|  | CI | 0.87 - 0.91 | 0.28 - 0.4 | 0.52 - 0.64 |
| | $r$ | 0.92*** | 0.6*** | 0.76*** |
| 28 | CCC | 0.93 | 0.56 | 0.75 |
|  | CI | 0.91 - 0.94 | 0.5 - 0.61 | 0.71 - 0.79 |
| | $r$ | 0.94*** | 0.82*** | 0.88*** |
| 35 | CCC | 0.94 | 0.70 | 0.86 |
|  | CI | 0.93 - 0.95 | 0.64 - 0.75 | 0.83 - 0.89 |
| | $r$ | 0.95*** | 0.83*** | 0.9*** |

Table 4 summarizes broad sense heritability (H^2^) values according to evaluation method, treatment and sampling day. Greater values of H^2^ (values closer to 1) were obtained using the digital images method. Similar results for H^2^ were obtained by other authors when comparing image based phenotyping methods to visual evaluations (Makanza et al. 2018; Singh et al. 2019). A phenotyping procedure (e.g., digital imaging) that detects high heritability of any given trait allows a broader selection process, hence, the genetic advance through the breeding cycles is faster (Holland et al. 2003).

**Table 4.**
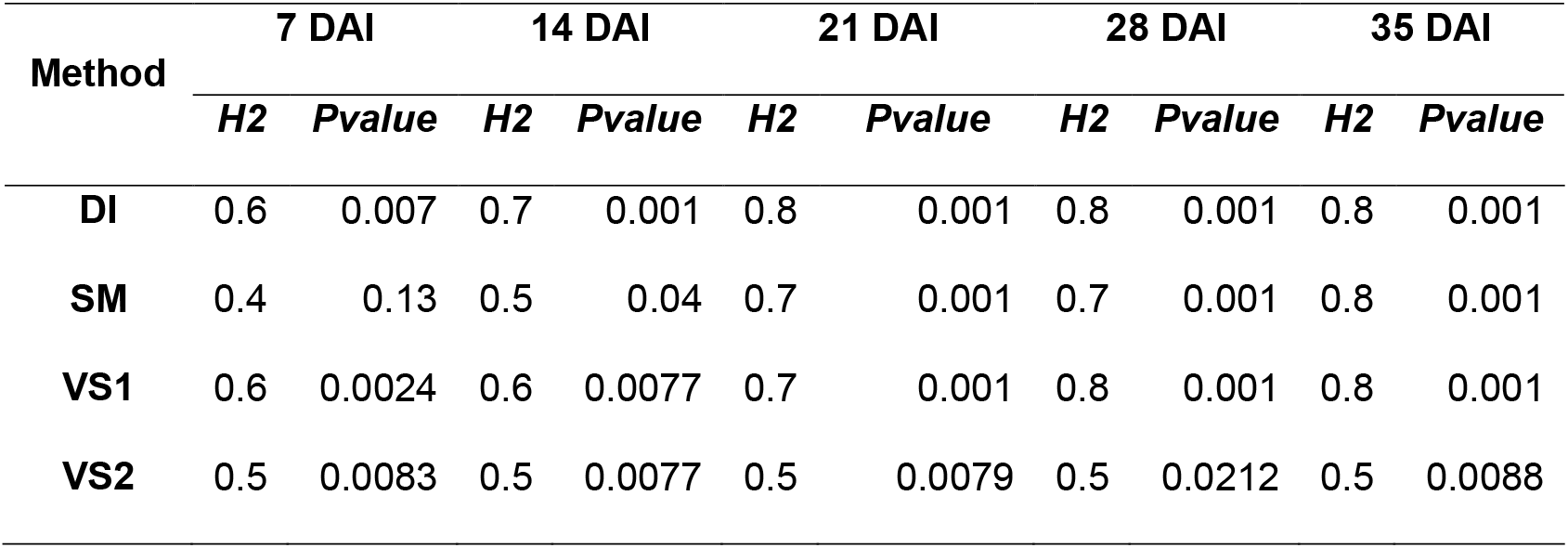
Broad sense heritability according treatment, evaluation method, and days after infestation for damage percentage. DI: Digital images; SM: SPAD measurements; VSI; Visual evaluation from expert; VS2: Visual evaluation from inexpert. DAI: Days after infestation.

| Method | 7 DAI |  | 14 DAI |  | 21 DAI |  | 28 DAI |  | 35 DAI |  |
| --- | --- | --- | --- | --- | --- | --- | --- | --- | --- | --- |
|  | *H2* | *Pvalue* | *H2* | *Pvalue* | *H2* | *Pvalue* | *H2* | *Pvalue* | *H2* | *Pvalue* |
| **DI** | 0.6 | 0.007 | 0.7 | 0.001 | 0.8 | 0.001 | 0.8 | 0.001 | 0.8 | 0.001 |
| **SM** | 0.4 | 0.13 | 0.5 | 0.04 | 0.7 | 0.001 | 0.7 | 0.001 | 0.8 | 0.001 |
| **VS1** | 0.6 | 0.0024 | 0.6 | 0.0077 | 0.7 | 0.001 | 0.8 | 0.001 | 0.8 | 0.001 |
| **VS2** | 0.5 | 0.0083 | 0.5 | 0.0077 | 0.5 | 0.0079 | 0.5 | 0.0212 | 0.5 | 0.0088 |

### Conclusions

The present work showed that estimation of plant damage from digital images yielded similar results to those obtained by the standard method of visual scoring by an expert evaluator. One of the major drawbacks of visual scoring is the dependence of an expert evaluator. Training of new evaluators for visual scoring of plant damage might be a straightforward mechanism to ensure continuity of such method over time. However, as argued before, inter-rater variation represents a major drawback for this method. Overall, SPAD measurements provided the less throughput and less agreement (CCC) and correlation with the standard evaluation of visual scoring from an expert, which makes this method an unattractive one to a breeding program. Higher values of broad sense heritability and faster recording of plant damage from digital images suggests that this phenotyping method might improve the efficiency of breeding for increased tolerance to spittlebugs in *U. humidicola*.

## Acknowledgments

We would like to thank William Mera, Ximena Bonilla, Jeison Velasco and Miller Escobar of the Tropical Forages Program (Alliance Bioversity-CIAT) for their technical support. This study was supported by the CGIAR Research Program 3.7 (Livestock and Fish).

## References

Araus JL, Kefauver SC, Zaman-Allah M, Olsen MS, Cairns JE. 2018. Translating High-Throughput Phenotyping into Genetic Gain [Internet]. [accessed 2020 Oct 8]. https://doi.org/10.1016/j.tplants.2018.02.001

Bock CH, Barbedo JGA, Del Ponte EM, Bohnenkamp D, Mahlein A-K. 2020. From visual estimates to fully automated sensor-based measurements of plant disease severity: status and challenges for improving accuracy. Phytopathol Res. 2(9).

Bock CH, Hotchkiss MW, Wood BW. 2016. Assessing disease severity: Accuracy and reliability of rater estimates in relation to number of diagrams in a standard area diagram set. Plant Pathol. 65(2):261–272.

Bock CH, Poole GH, Parker PE, Gottwald TR. 2010. Plant disease severity estimated visually, by digital photography and image analysis, and by hyperspectral imaging. CRC Crit Rev Plant Sci. 29:59–107.

Büchi L, Wendling M, Mouly P, Charles R. 2018. Comparison of Visual Assessment and Digital Image Analysis for Canopy Cover Estimation. Agron J [Internet]. [accessed 2020 Oct 8] 110(4):1289–1295. http://doi.wiley.com/10.2134/agronj2017.11.0679

Cardona C, Miles JW, Sotelo G. 1999. An Improved Methodology for Massive Screening of *Brachiaria* spp. Genotypes for Resistance to *Aeneolamia varia* (Homoptera: Cercopidae). J Econ Entomol [Internet]. [accessed 2019 May 21] 92(2):490–496. http://ciat-library.ciat.cgiar.org/Articulos_Ciat/jee.pdf

Cardoso JA, Rao IM. 2019. Drought Resistance of Tropical Forage Grasses. In: Pessarakli M, editor. Handb Plant Crop Stress [Internet]. 4th ed. Boca Raton: CRC Press; [accessed 2020 Oct 8]; p. 793–803. https://www.taylorfrancis.com/

Cardoso JA, Rincon J, Jimenez J de la C, Noguera D, Rao IM, Rincón J, De La Cruz Jimenez J, Noguera D, Rao IM. 2013. Morpho-anatomical adaptations to waterlogging by germplasm accessions in a tropical forage grass. AoB Plants. 5:plt047.

Holland JJB, Nyquist WWE, Cervantes-Martínez CT. 2003. Estimating and Interpreting Heritability for Plant Breeding: An Update. Plant Breed Rev [Internet]. [accessed 2020 Oct 8] 22(December):9–112. http://doi.wiley.com/10.1002/9780470650202.ch2

El Jarroudi M, Kouadio AL, Mackels C, Tychon B, Delfosse P, Bock CH. 2015. A comparison between visual estimates and image analysis measurements to determine septoria leaf blotch severity in winter wheat. Plant Pathol [Internet]. [accessed 2020 Oct 8] 64(2):355–364. http://doi.wiley.com/10.1111/ppa.12252

Jiménez J de la C, Cardoso JA, Leiva LF, Gil J, Forero MG, Worthington ML, Miles JW, Rao IM. 2017. Non-destructive Phenotyping to Identify Brachiaria Hybrids Tolerant to Waterlogging Stress under Field Conditions. Front Plant Sci [Internet]. [accessed 2020 Mar 23] 8(167). www.frontiersin.org

Jiménez J de la C, Leiva L, Cardoso JA, French AN, Thorp KR. 2020. Proximal sensing of Urochloa grasses increases selection accuracy. Crop Pasture Sci [Internet]. [accessed 2020 Oct 8] 71(4):401–409. http://www.publish.csiro.au/?paper=CP19324

Makanza R, Zaman-Allah M, Cairns JE, Magorokosho C, Tarekegne A, Olsen M, Prasanna BM. 2018. High-throughput phenotyping of canopy cover and senescence in maize field trials using aerial digital canopy imaging. Remote Sens. 10(2).

Mazabel J, Worthington M, Castiblanco V, Peters M, Arango J. 2020. Using near infrared reflectance spectroscopy for estimating nutritional quality of Brachiaria humidicola in breeding selections. Agrosystems, Geosci Environ [Internet]. [accessed 2020 Oct 8] 3(e20070). https://onlinelibrary.wiley.com/doi/10.1002/agg2.20070

Miles JW, Cardona C, Sotelo G. 2006. Recurrent selection in a synthetic brachiariagrass population improves resistance to three spittlebug species. Crop Sci. 46(3):1088–1093.

Nita M, Ellis MA, Madden L V. 2003. Reliability and accuracy of visual estimation of phomopsis leaf blight of strawberry. Phytopathology. 93(8):995–1005.

Parsa S, Sotelo G, Cardona C. 2011. Characterizing herbivore resistance mechanisms: Spittlebugs on Brachiaria spp. as an example. J Vis Exp [Internet].(52). http://www.jove.com/details.php?id=3047

Piepho H-P, Möhring J. 2007. Computing heritability and selection response from unbalanced plant breeding trials. Genetics. 177:1881–1888.

R Development CoreTeam. 2015. A language and environment for statistical computing.

Shakoor N, Lee S, Mockler TC. 2017. High throughput phenotyping to accelerate crop breeding and monitoring of diseases in the field. Curr Opin Plant Biol. 38:184–192.

Singh D, Wang X, Kumar U, Gao L, Noor M, Imtiaz M, Singh RP, Poland J. 2019. High-throughput phenotyping enabled genetic dissection of crop lodging in wheat. Front Plant Sci. 10(394).

Valério J.., Cardona C, Peck D.., Sotelo G. 2001. Spittlebugs: bioecology, host plant resistance and advances in IPM. In: Proc XIX Int Grassl Congr Piracicaba, FEALQ [Internet]. Sao Pedro, Piracicaba; [accessed 2019 Jul 16]; p. 217–221. https://cgspace.cgiar.org/bitstream/handle/10568/56032/tema5-2.pdf?sequence=1&isAllowed=y

Vasques Berchembrock Y, de Figueiredo UJ, Rodrigues Nunes JA, Borges do Valle CB, Lima Barrios SC. 2020. Comparison of selection methods among and within full-sibling progenies in Urochloa humidicola. Grass Forage Sci. 75(2):145–152.

Walter A, Studer B, Kölliker R. 2012. Advanced phenotyping offers opportunities for improved breeding of forage and turf species. Ann Bot. 110:1271–1279.

